# Comparative analysis of dietary iron deprivation and supplementation in a murine model of colitis

**DOI:** 10.1101/2024.12.19.629342

**Authors:** Thanina Medjbeur, Ugo Sardo, Prunelle Perrier, Kevin Cormier, Marilyne Roy, Anne Dumay, Léon Kautz

## Abstract

Inflammatory bowel diseases are chronic inflammatory conditions with growing prevalence in western populations. Iron is an essential component of erythrocytes hemoglobin. Under the influence of elevated hepcidin production, iron is sequestered in cells during inflammation which, in turn, leads to iron restriction for red blood cells synthesis. As a consequence, iron deficiency and anemia of inflammation are the most prevalent extraintestinal complications in IBD patients. Patients are thus treated with oral iron supplements that have limited efficacy as iron absorption is blunted during intestinal inflammation. Moreover, iron supplementation can cause intestinal complications and previous studies have shown that iron supplementation worsens the inflammatory response. However, a comparative analysis of the effects of low, adequate and high dietary iron content matching iron the supplementation given to patients has not been performed in mice. We therefore tested the impact of dietary iron deprivation and supplementation in a murine model of colitis induced by dextran sodium sulfate. We found that both dietary iron deprivation and supplementation were accompanied by a more severe inflammation with earlier signs of gastrointestinal bleeding compared to mice fed an iron adequate diet. The manipulation of dietary iron led to a comparable oxidative stress and a pronounced dysbiosis in the colon of control mice that differed depending on the dietary iron content. Analysis of these dysbiosis is in line with a pronounced susceptibility to colonic inflammation thus questioning the benefit/risk balance of oral iron supplementation for IBD patients.

## INTRODUCTION

Iron is a trace element essential for nearly all living species but excess iron catalyzes the generation of reactive oxygen species. In humans, most of the plasma iron destined for erythropoiesis is provided by the recycling of iron from senescent red blood cells by splenic macrophages and 25% of the body iron is stored in hepatocytes(1). Iron losses are minor and compensated by the absorption of dietary iron by duodenal enterocytes. Iron is released into the circulation by the exporter ferroportin present at the cell surface of macrophages, enterocytes and hepatocytes. This process is orchestrated by the liver-produced hormone hepcidin(2), which binds to ferroportin and induces its occlusion and degradation leading to decreased iron uptake. In conditions associated with chronic inflammation, increased hepcidin production(3) prevents iron efflux into the bloodstream and causes a profound iron restriction that paves the way for the development of anemia, a frequent condition in hospitalized or chronically ill patients.

Ulcerative colitis and Crohn’s disease, collectively termed inflammatory bowel diseases (IBD), are chronic forms of intestinal inflammation affecting the ileum and the large intestine of more than 300/100.000 individuals in Western populations(4). Symptoms include abdominal pain, rectal bleeding, diarrhea, weight loss and nutritional deficiencies. Since iron deficiency and anemia of inflammation are the most prevalent extraintestinal complication(5), IBD patients are commonly treated with oral iron supplements that can cause abdominal pain and gastrointestinal complications(6–8) and increase ROS production and the risk of carcinogenesis(9). Indeed, oral iron supplementation in IBD patients and increased dietary iron concentration in murine models of colitis are associated with exacerbated intestinal inflammation, blunted immune response(9) and colonic dysbiosis(10, 11). In contrast, mice fed an iron-deficient diet seemed less susceptible to intestinal inflammation during experimental colitis(12). In most studies, a standard rodent chow was used to study the effect of dietary iron but a concomitant comparison of the outcome of low and high dietary iron content mimicking oral iron supplementation during colitis has not been performed. Here, we sought to compare the impact of dietary iron deprivation and supplementation on intestinal inflammation in a mouse model of colitis induced by dextran sulfate sodium.

## MATERIAL AND METHODS

### Animals

5-week-old C57Bl/6J male mice (n=48) were purchased at Janvier laboratories (Le Genest St-Isle, France) and housed in the CREFRE US006 animal care facility. Custom made diets of identical composition (Ssniff) were supplemented (50 and 8000µg/g) or not (10µg/g) with carbonyl iron. At 6 weeks of age, mice were fed either an iron deficient (ID, 10µg/g Fe) or an iron adequate (IA; 50µg/g Fe) diet for 2 weeks, with access to water ad libitum. After two weeks, a group of mice fed an IA diet was given an iron-enriched diet (IE; 8000µg/g Fe) on the day the colitis protocol was initiated. The iron content of the iron-enriched diet was chosen based on the physiological iron needs in humans and rodents. Every day, a healthy 70 kg adult human needs to absorb 1-2 mg of iron from the diet which equals to 0,03 mg/kg. Mice require the daily intake of 10µg iron which, on the basis of a 20g mouse, amounts to 1 mg/kg, approximately 33-times more than humans. Patients are usually supplemented with 100-200 mg oral iron supplements daily thus exceeding the iron needs by 100-fold which would translate to an intake of 1000µg in mice. We applied the correction factors of mice vs human needs (x33) and decided to supplement the diet to reach an intake of 32,000µg Fe/day. As mice consume 4 grams of food a day, our custom-made IE diet therefore contains 8000µg/g of iron. All mice were caged in a specific pathogen-free animal facility with controlled temperature, humidity and dark-light cycles of 12 hours each. Samples were harvested, immediately frozen in liquid nitrogen and stored at -80°C. Procedures were approved by the Animal Care and Ethics Committee of US006/CREFRE (CEEA-122; application number APAFIS 2016092912382223)

### Experimental colitis

Mice were administered 2% Dextran Sodium Sulfate (DSS, MP Biomedicals) in drinking water for 7 days and switched to regular drinking water. Animals were monitored daily for signs of illness and/or welfare impairment and were euthanized by cervical dislocation on day 9. Body weight and disease activity index (DAI) calculated using the scoring system shown in Table 1A, were recorded every day.

**Table 1:** Assessment of inflammatory phenotype. (A) Disease activity index (DAI) scoring chart. (B) Histological microscopic scores chart.

| Score | 0 | 1 | 2 | 3 |
| --- | --- | --- | --- | --- |
| Weight loss (%) | 0 | 0 to 10 | 10 to 20 | >20 |
| Stool consistency | Normal | Loose | Diarrhea | - |
| Blood in the feces | No | Yes | - | - |
| Mouse appearance | Normal | Prostrate | Prostrate with spiky hairs | Lethargic |
| Maximal score | 9 |  |  |  |

| Score | 0 | 1 | 2 | 3 |
| --- | --- | --- | --- | --- |
| Submucosal oedema | Absent | Mild | Moderate | Severe |
| Vasculitis | No | Yes | - | - |
| Mucosal architecture | Conserved | Mild | Moderate | Lost |
| Cell infiltration | No | Mild | Moderate | Severe |
| Muscle thickening | No | Yes | - | - |
| Crypt abscesses | No | Yes | - | - |
| Goblet cell depletion | No | Yes | - | - |
| Maximal score | 13 |  |  |  |

### Histological assessment of colonic injury

Colonic tissue specimens were excised 2 cm proximal to the anus and immediately transferred into 4% formaldehyde before paraffin embedding. Five-micrometer colonic sections were stained with hematoxylin-eosin. Histological damage was evaluated based on inflammatory cell infiltration, epithelial/mucosal alteration (including vasculitis, goblet cell depletion, and crypt abscesses), mucosal architecture alteration (including ulceration and crypt loss), and submucosal oedema (Table 1B). Colonic sections were incubated in 1% periodic acid (Sigma, 375810) for 15 minutes, in Schiff reagent (Sigma, 320680) for 15 minutes and counterstained with Mayer’s hematoxylin (RAL DIAGNOSTICS, 361075) to stain for polysaccharides, glycoproteins, and acidic mucopolysaccharides.

### Iron parameters

Serum iron concentration and unsaturated iron binding capacity were determined using iron direct and UIBC kits (Biolabo) according to the manufacturer’s instructions. Liver iron content and iron deposition in the colon (Perl’s stain) were determined as previously described(13).

### Gene expression

Total RNA was isolated using Trizol-Chloroform. 10 µg of RNA were purified using the Dynabeads mRNA Direct kit to eliminate residual DSS. Complementary DNA was synthesized using MMLV reverse transcriptase (Promega). Real-time qPCR reactions were prepared with Takyon SYBR Mastermix (Eurogentec) and run on a LightCycler 480 System (Roche Diagnostics). The following forward (F) and reverse (R) primers were used:

*Hprt: F-* CTGGTTAAGCAGTACAGCCCCAA, R-CGACAGGTCCTTTTCACCAGC; *Hamp*: F-AAGCAGGGCAGACATTGCGAT, R-CAGGATGTGGCTCTAGGCTATGT; *Tnf*α: F-AATGGCCTCCCTCTCATCAG, R-GCTACGACGTGGGCTACAGG; *Inf*γ: F-CAGCAACAGCAAGGCGAAA, R-AGCTCATTGAATGCTTGGCG; *Il6*: F-CTCTGCAAGAGACTTCCATCCAGT, R-CGTGGTTGTCACCAGCATCA; *Il1*β: F-ACCTTCCAGGATGAGGACATGAG, R-CATCCCATGAGTCACAGAGGATG; *Il1*0: F-AGGCGCTGTCATCGATTTCTC, R-TGCTCCACTGCCTTGCTCTTA; *Il17a*: F-TCCAGAAGGCCCTCAGACTA, R-CAGGATCTCTTGCTGGATG; *Il22*: F-AGGTGGTGCCTTTCCTGACC, R-ACCGCTGATGTGACAGGAGC; *Cxcl9*: F-GGCAAATGTGAAGAAGCTGATG, R-TTTTTCCCCCTCTTTTGCTTTT; *Cxcl10*: F-TGTTGAGATCATTGCCACGA, R-CCAGTTAAGGAGCCCTTTTAGACC; *Reg3*β: F-TGGTTTGATGCAGAACTGGC, R-TGGAGGACAAGAATGAAGCCT; *Reg3*γ: F-CCTCCATGATCAAAAGCAGTGG, R-GGATTCGTCTCCCAGTTGATGT; *Ocln*: F-TGGATGACTACAGAGAGGAGAGT, R-TCCTCTTGATGTGCGATAATTTGC; *Tjp1*: F-CACAGCCTCCAGAGTTTGACAG, R-TCCACAGCTGAAGGACTCACAG; *Cldn2*: F-TCTCAGCCCTGTTTTCTTTGGT, R-GGGCCTGGTAGCCATCATAGTA; *Cldn8*: F-GGAGGAGCACTGTTCTGTTGTG, R-GTGGAAACTCCGTTGAGTGGT; *Muc2*: F-TGTCCCGACTTCAACCCAAG, R-TCTGGTTTTGAGGGATGCATGT; *Muc4*: F-AGAGGCAGAAGAGGAGTGGAGA, R-GGTGGTAGCCTTTGTAGCCATC; *Ptgs2*: F-GCCTCCCACTCCAGACTAGA, R-ACAGCTCAGTTGAACGCCTT; *Chac1*: F-AAGATGAGCACCTGGAAGCC, R-CTTGGCTCCTCAGGTCAGTG; *Ascl4*: F-TGGTCAGGGATATGGGCTGA, R-CCACCTTCCTGCCAGTCTTT; *Sod1*: F-AGGAGAGCATTCCATCATTGG, R-CCCAGCATTTCCAGTCTTTGT; *Sod2*: F-GGGCTGGCTTGGCTTCAATAA, R-TCCCACACGTCAATCCCCAG; *Lcn2*: F-TCTGTCCCCACCGACCAAT, R-CCAGTCAGCCACACTCACCAC; *Hmox1*: F-CAGATGGCGTCACTTCGTCA, R-CTCTGCAGGGGCAGTATCTTG; *Nqo1*: F-AGCGGCTCCATGTACTCTCT, R-GCATCTGGTGGAGTGTGGC; *Atg5:* F-CAACCGGAAACTCATGGAAT, R-CGGAACAGCTTCTGGATGA; *Atg7*: F-GGTCTTACCCTGCTCCATCA, R-TGTGGTTGCTTGCTTCAGAG; *Xbp1s*: F-GAACATCTTCCCATGGACTC, R-CCCAAAAGGATATCAGACTCAG.

The target gene expression was normalized to the Hypoxanthine Phosphoribosyl Transferase (*Hprt*) and quantified using the comparative -ΔCt (threshold cycle) method.

### Microbiota analysis by 16S rDNA sequencing

DNA was extracted from fecal samples by using the QIAamp PowerFecalPro DNA kit (Qiagen, Hilden,MD) according to the manufacturer’s recommendations. DNA extracts were quantified on a Qubit4 fluorometer using the dsDNA HS Assay Kit (Life Technologies; USA). For 16S rDNA gene sequencing, a two-step PCR library preparation was applied, according to the recommendations of the 16S Metagenomic Sequencing Library Preparation Guide (Illumina, San Dego, CA, USA). Briefly, for the first PCR reaction, V4 hyper-variable region of the 16S rDNA were amplified using the primers 515-F: 5’- *TCG TCG GCA GCG TCA GAT GTG TAT AAG AGA CAG **G*****TG CCA GCM GCC GCG GTAA -**3’ and 806-R: 5’- *GTC TCG TGG GCT CGG AGA TGT GTA TAA GAG ACA G***GG ACT ACH VGG GTW TCT AAT-**3’ (italicized sequence is the Illumina adapter following by barcode, bold sequence is the conserved bacterial sequence). Amplicons were then purified using AMPure XP beads (Beckman Coulter, Indianapolis, IN). A second PCR reaction was performed to incorporate sample-specific barcode, using Nextera XT index kit (Illumina, USA). After amplicons purification, DNA concentration was controlled by qPCR using KAPA Library Quantification kit (Roche). A master DNA pool was then generated in equimolar ratios. PhiX Control v3 (Illumina, USA) was added to check quality of run. The pooled products quantity were controlled on Qubit4 fluorometer, load into an Illumina MiSeq cartridge and sequenced (paired-end reads, 2 × 300 bp) on IlluminaMiSeq sequencer. At the end of the run, FastQ files were generated.

Sequences were demultiplexed, reads quality control was done and paired-end amplicon reads were processed using the FROGS pipeline (Find Rapidly OTU with Galaxy Solution, Version 4.1.0) on the Galaxy Migale platform (https://galaxy.migale.inra.fr/) (14). Briefly, forward and reverse reads were trimmed for adaptor and PCR primers removal, merged, and chimeric sequences were removed. Reads were then clustered in Amplicon Sequence Variants (ASVs) with an aggregation distance of 1, and filtered with a minimum relative abundance threshold of 0.005% and blast coverage > 0.95. Taxonomic assignment was performed against the 16S SILVA 138 pintail 100 database. Before futher analysis, all samples were rarefied to the same depth, with a minimum reads number of 74,959. Bacterial composition and diversities were estimated using FROGSTAT phyloseq tools. Alpha-diversity within group was estimated by calculating richness (Observed) and Shannon diversity index. Beta-diversity between groups was evaluated by calculating Bray-Curtis distances between samples. Ordination using principal coordinates analysis (PcoA) was performed to represent biodiversity distribution at the ASV level between groups. To identify differentially abundant taxa, DESeq2(15) was used.

### Statistical analysis

The statistical significances were assessed by Student’s t-test, one or two-way analysis of variance (ANOVA) specified in the figure legend using Prism 10 (GraphPad). For bacterial abundance and alpha-diversity, significant differences were analyzed by ANOVA with Bonferroni multiple comparison test. The significance of beta-diversity metrics was assessed by PERMANOVA test (Permutational Multivariate Analysis of Variance Using Distance Matrices).

## RESULTS

### Iron metabolism during DSS-induced colitis

6 weeks old C57BL/6 male mice were switched from a standard rodent chow to an iron-adequate (IA) or iron-deficient (ID) diet for 2 weeks (Figure 1A). A third group was fed an IA diet for 2 weeks and switched to an iron-enriched diet (IE) when the colitis was initiated in order to mimic the supplementation given to patients. Mice were then administered 2% DSS in drinking water for 7 days and returned to regular water until the peak inflammation was reached which, according to our institutional regulations, required euthanasia by day 9. Control mice for each diet were given water for 9 days. Histological assessment of non-heme iron distribution in the colon by Prussian blue staining showed that iron accumulated in the lumen and enterocytes of control mice fed an IE diet whereas no iron was detectable in the colon of mice fed an ID or IA diet (Figure 1B). A milder accumulation of iron was observed in the colon of DSS-treated mice fed an IE diet (Figure 1C). Serum iron concentration and transferrin saturation decreased significantly in DSS-treated mice compared to control mice regardless of the diet (Figure 1D). After DSS, mice fed an ID diet had slightly lower transferrin saturation compared to mice fed an IA diet but plasma iron levels were similar between mice fed an IA or IE diet. While liver iron concentration decreased in DSS-treated mice fed an ID or IA diet compared to their respective controls, liver iron content was increased in mice fed an IE diet. These results indicate that iron supplementation was only moderately effective at mitigating the iron deficiency caused by colonic inflammation.

**Figure 1:**
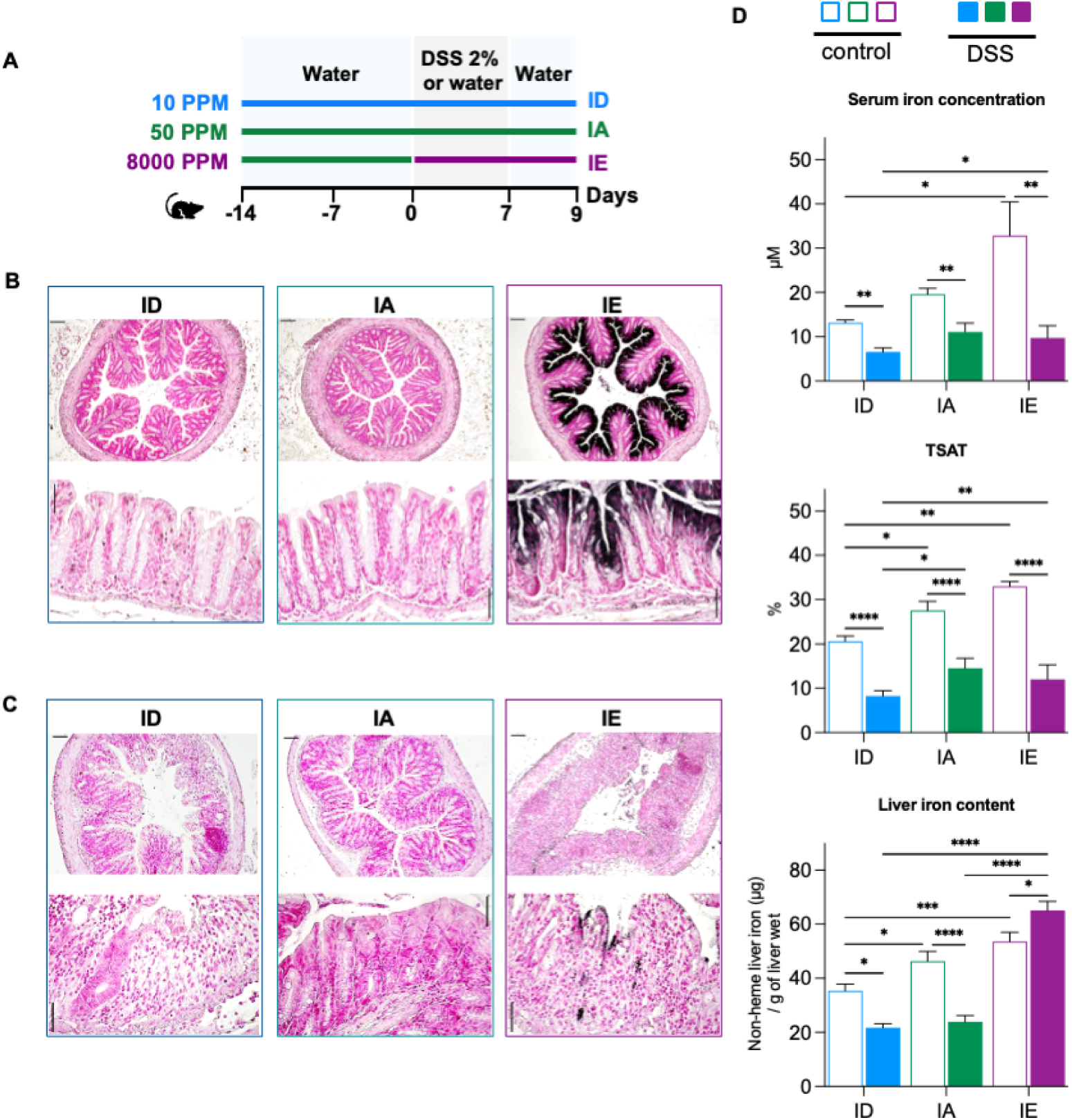
Iron metabolism during DSS-induced colitis. (A) Mice fed an iron deficient (ID), adequate (IA), or enriched (IE) diet were given regular drinking water (control; empty bars) or 2% dextran sodium sulfate (DSS) for 7 days and regular drinking water for 2 days (DSS; plain bars). Iron deposition was assessed by Perls’ staining on colon tissue section from control (B) or DSS-treated mice (C). Upper row scale bar = 100µm; lower row scale bar = 50µm. (D) Serum iron concentration, transferrin saturation and liver iron content in control and DSS-treated mice. Data shown are means ± s.e.m and were compared between DSS-treated mice and control mice for each diet (n = 8) by Two-way ANOVA and corrected for multiple comparisons by Holm-Šidák method. ****P < 0.0001, ***P < 0.001, **P < 0.01, *P < 0.05.

### Iron deprivation and supplementation worsen the inflammatory phenotype

Control mice did not show any significant difference in body weight (Figure 2A) or disease activity index (Figure 2B). Only a trend toward a decrease in body weight that was rapidly compensated was observed in mice fed an IE diet, presumably as a result of the change in diet. However, mice fed an ID and IE diet exhibited a more pronounced weight loss than mice fed an IA diet 6 to 9 days after DSS administration (Figure 2A) with an average body weight loss of 20%. The weight loss was accompanied by a more severe disease activity index in mice fed an ID or IE diet compared to mice fed an IA diet, with a significant increase as early as 3-4 days after DSS administration (Figure 2B). The presence of blood was detected in the cage of mice fed an ID diet 1 day after DSS administration (Figure 2C) and the decrease in stool consistency was observed after 2-3 days in mice fed an ID or IE diet, 2 days before the first signs in mice fed an IA diet (Figure 2D). The change in overall mouse appearance was also more pronounced in mice fed an ID and IE diet compared to mice fed an IA diet (Figure 2E). While the microscopic scores increased in all the mice subjected to DSS compared to their respective controls, mice fed an IE diet showed more severe inflammatory scores compared to mice fed an IA diet (Figure 2F-G). Altogether, these results indicate that mice fed an ID or IE diet exhibit a more severe colonic inflammation compared to mice fed an IA diet.

**Figure 2:**
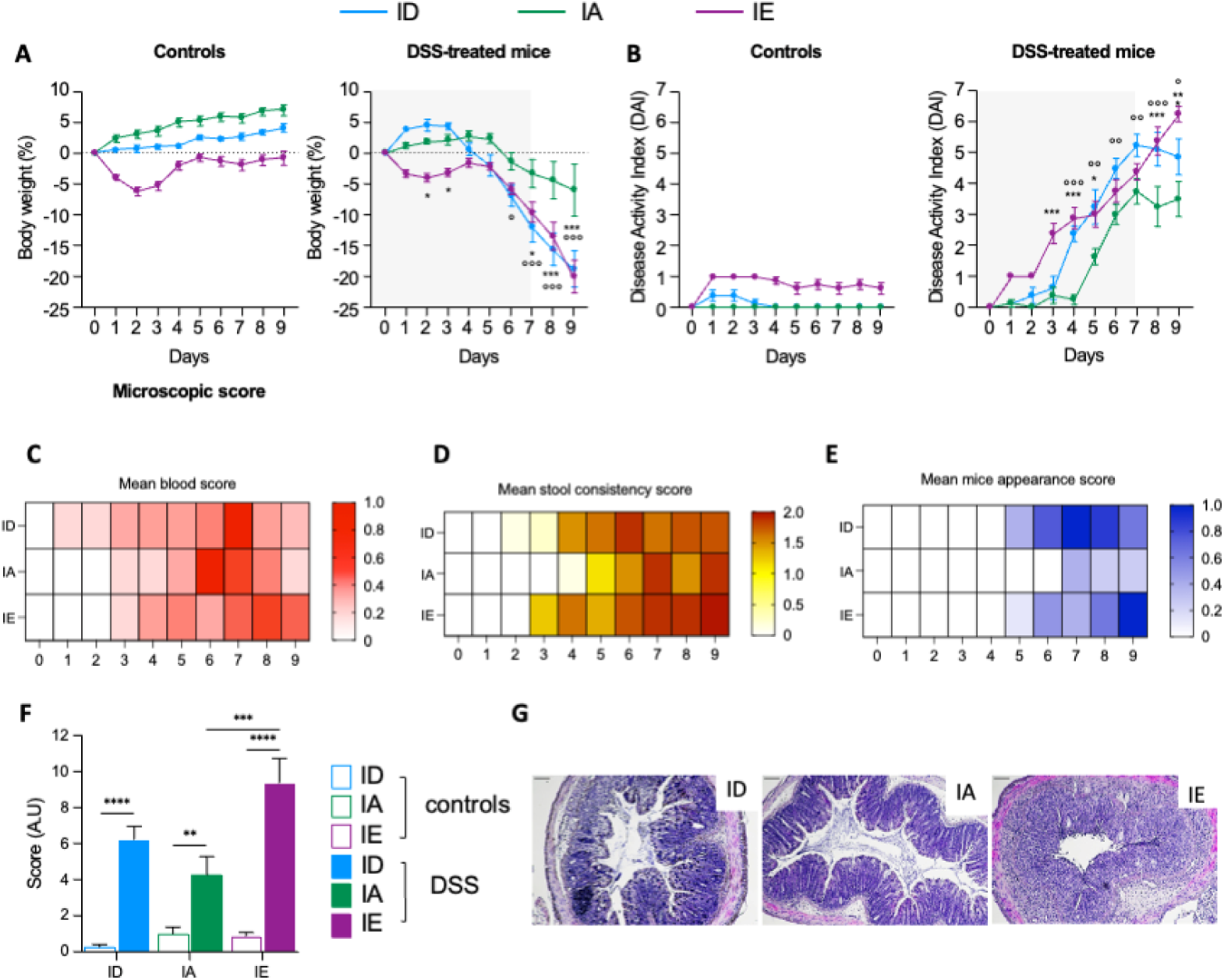
Iron deprivation and supplementation worsen the inflammatory phenotype during DSS-induced colitis. (A) Body weight (%), (B) disease activity index (DAI), (C) mean blood score, (D) mean stool consistency score, (E) mean appearance score and (F) histological microscopic score in control and DSS-treated mice. (G) hematoxylin-eosin staining of colon sections from DSS-treated mice. Scale bar = 50µm. Data shown are means ± s.e.m and were compared between DSS-treated mice and control mice for each diet (n = 8) by Two-way ANOVA and corrected for multiple comparisons by Holm-Šidák method. (A-B) IE vs IA: ***P < 0.001, **P < 0.01, *P < 0.05; ID vs IA: °°°P < 0.001, °°P < 0.01, °P < 0.05. (F) ****P < 0.0001, ***P < 0.001, **P < 0.01, *P < 0.05.

### Transcriptional response to dietary iron challenge during colonic inflammation

Consistent with an alteration of epithelial barrier integrity during DSS induced colitis(16), tight junction proteins Occludin 1 (*Ocln*) and tight junction protein 1 (*Tjp1*) mRNA expression was significantly reduced 9 days after DSS administration in the colon of mice fed an ID diet (Figure 3A). A comparable decrease in *Ocln* expression in mice fed an IE diet was accompanied by a decrease in claudin 2 and 8 (*Cldn2*, *Cldn8*) expression. The dietary iron content did not induce any change in *Ocln*, *Tjp1*, *Cldn2*, *Cldn8* expression in DSS-treated mice (Figure 3B). Expression of mucin 2 (*Muc2*) and 4 (*Muc 4*) was increased in mice fed an IE diet compared to mice fed an IA diet (Figure 3A-B). However, histological assessment of colon section by periodic acid–Schiff (PAS) staining indicated that the number of goblet cells per crypt was reduced upon DSS treatment and to a greater extent in mice fed an ID or IE diet compared to mice fed an IA diet (Figure 3C). As epithelial damage is usually associated with mucosal inflammation, we next assessed whether dietary iron influenced the expression level of inflammatory cytokines (*Tnf*α*, Ifn*γ*, Il17a, Il1*β*, Il6, Il10* and *Il22*), chemokines (*Cxcl9* and *Cxcl10*), and antimicrobial peptides (*Reg3*β*, Reg3*γ) known to contribute to the inflammatory response in the colon (Figure 4A-B). Expression of *Tnf*α*, Il17a, Il1*β, and *Il6* was increased in DSS-treated mice fed an ID or IE diet compared to their respective controls whereas expression of chemokines *Cxcl9* and *Cxcl10* and antimicrobial peptides *Reg3*β*, Reg3*γ was markedly increased only in mice fed an IE diet and subjected to DSS-induced colitis (Figure 4A). A significant increase in *Tnf*α*, Il17a* and *Cxcl9* expression was observed in DSS-treated mice fed an IE diet compared to mice fed an IA diet (Figure 4B) suggesting that iron supplementation fueled the immune response. Previous studies have shown that ferroptosis, oxidative stress, autophagy and ER stress contribute to colonic inflammation during IBD(17). We therefore assessed whether iron deprivation or supplementation altered the expression level of mediators of ferroptosis (*Ptgs2, Acsl4, Chac1*), oxidative stress (*Sod1, Sod2, Lcn2, Hmox, Nqo1*), autophagy (*Atg5, Atg7*), ER stress (*Xbp1s*) in DSS-treated and control mice. DSS treatment only led to an increase in lipocalin 2 (*Lcn2*) expression regardless of the dietary iron content), which is consistent with its proposed function as an antimicrobial protein(18), and expression of the other markers was unchanged by inflammation (Figure 5A). However, after DSS treatment, expression of *Acsl4, Chac1*, *Sod1, Sod2* and *Nqo1* was significantly increased in the colon of mice fed an ID or IE diet compared to mice fed an IA diet suggesting that ferroptosis and oxidative stress are triggered upon iron deprivation and supplementation (Figure 5B).

**Figure 3:**
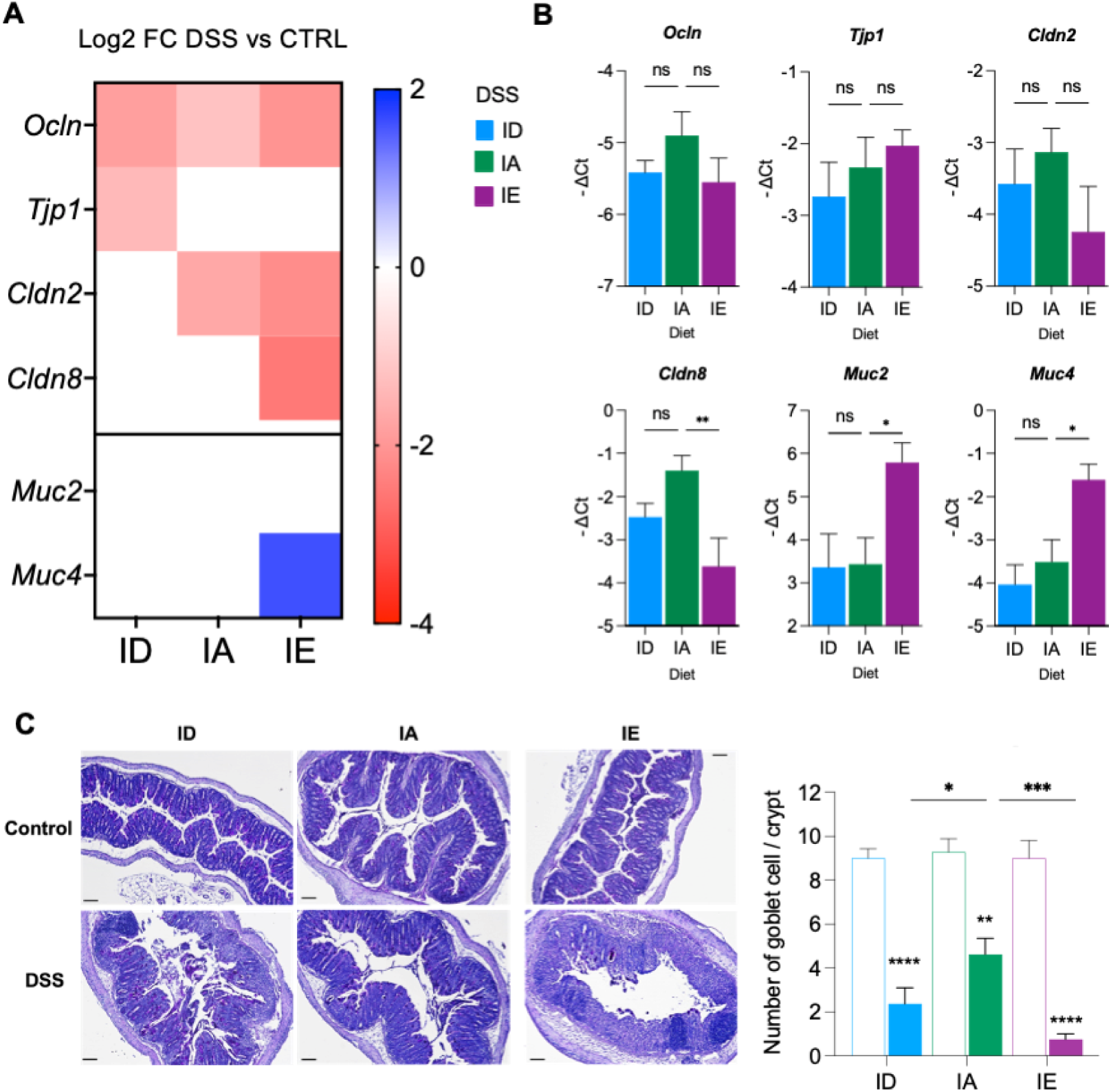
Intestinal barrier during colitis. (A) Representative heatmap of tight junction proteins Occludin 1 (*Ocln*), tight junction protein 1 / Zona occludens 1 (*Tjp1*), claudin 2 and 8 (*Cldn2*, *Cldn8*) and mucin 2 (*Muc2*) and 4 (*Muc 4*) mRNA expression in the colon of DSS-treated mice compared to their respective controls. Only differences reaching statistical significance (P<0.05) are represented. (B) mRNA expression in the colon of DSS-treated mice fed an ID or IE diet compared to mice fed an IA diet. (C) Periodic acid Schiff staining of colon section and the observed number of goblet cells per crypt. Scale bar = 50µm. Data shown are means ± s.e.m and were compared between DSS-treated mice and control mice for each diet (n = 8) by Two-way ANOVA and corrected for multiple comparisons by Holm-Šidák method. ****P < 0.0001, ***P < 0.001, **P < 0.01, *P < 0.05.

**Figure 4:**
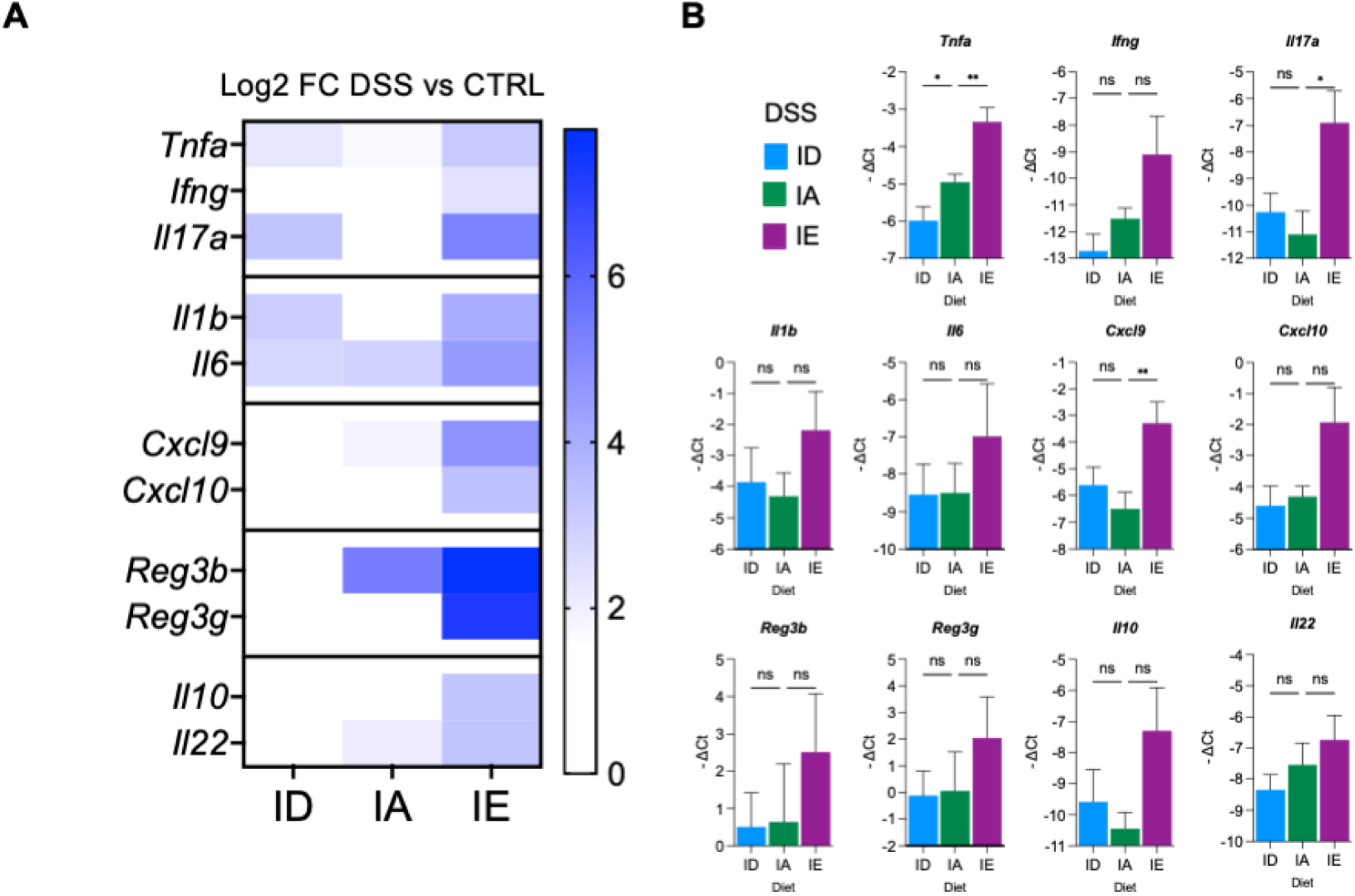
Expression of inflammatory and immune mediators during colitis. mRNA expression of inflammatory cytokines (*Tnf*α*, Ifn*γ*, Il17a, Il1*β*, Il6, Il10* and *Il22*), chemokines (*Cxcl9* and *Cxcl10*), and antimicrobial peptides (*Reg3*β*, Reg3*γ) in the colon of control and DSS-terated mice. (A) Representative heatmap of expression level in DSS-treated mice compared their respective controls. Only differences reaching statistical significance (P<0.05) are represented. (B) Expression level in DSS-treated mice fed an ID or IE diet compared to mice fed an IA diet. Data shown are means ± s.e.m and were compared between DSS-treated mice and control mice for each diet (n = 8) by Two-way ANOVA and corrected for multiple comparisons by Holm-Šidák method. **P < 0.01, *P < 0.05.

**Figure 5:**
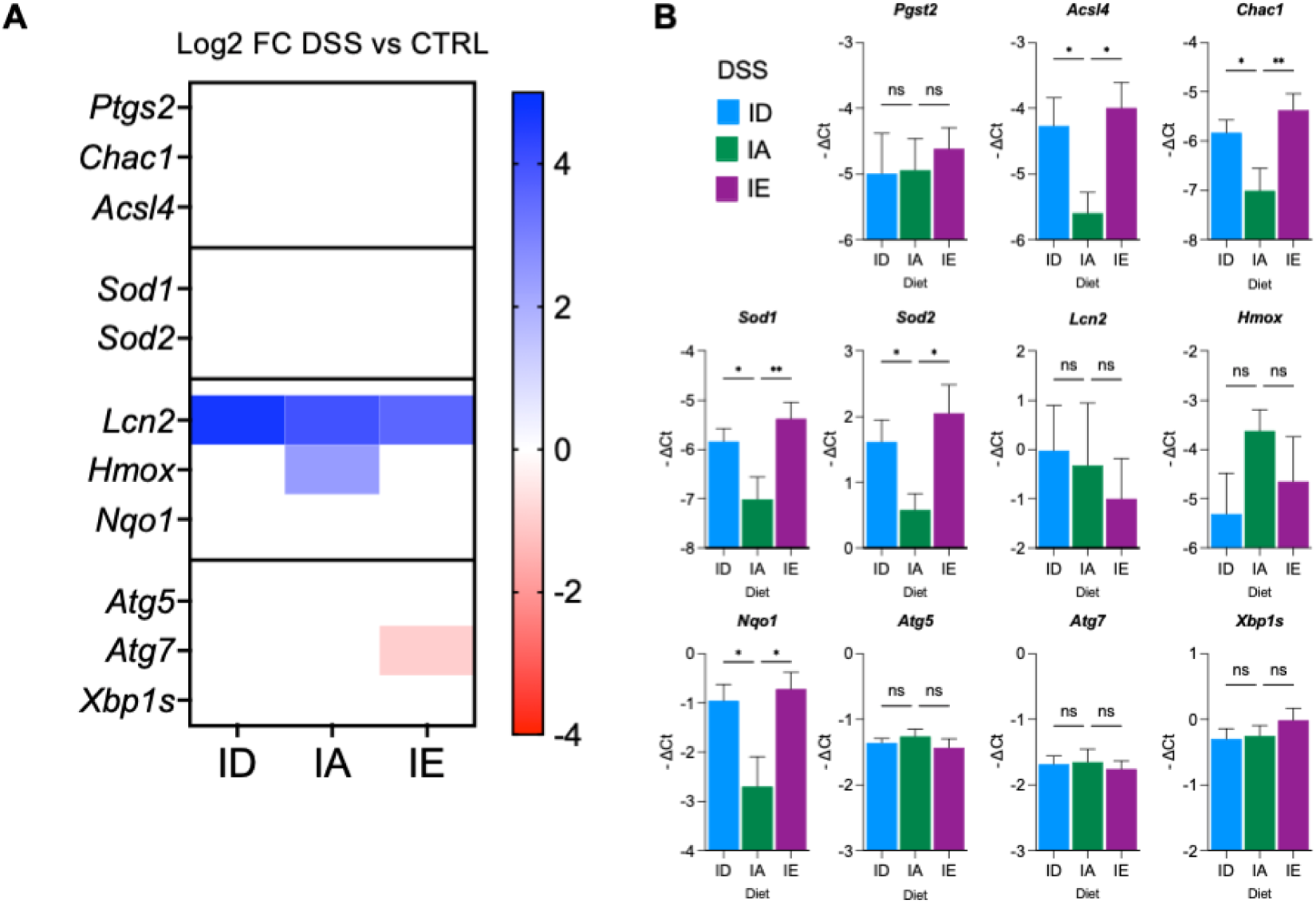
Dietary iron deprivation and supplementation induce oxidative stress and ferroptosis in DSS-treated mice. mRNA expression level of mediators of ferroptosis (*Ptgs2, Acsl4, Chac1*), oxidative stress (*Sod1, Sod2, Lcn2, Hmox, Nqo1*), autophagy (*Atg5, Atg7*), ER stress (*Xbp1s*) in the colon of control and DSS-treated mice. (A) Representative heatmap of expression level in DSS-treated mice compared to their respective controls. Only differences reaching statistical significance (P<0.05) are represented. (B) Expression level in DSS-treated mice fed an ID or IE diet compared to mice fed an IA diet. Data shown are means ± s.e.m and were compared between DSS-treated mice and control mice for each diet (n = 8) by Two-way ANOVA and corrected for multiple comparisons by Holm-Šidák method. **P < 0.01, *P < 0.05.

### Dietary iron impact on fecal microbiota diversity

We next investigated the impact of the dietary iron content on bacterial abundance and composition by 16S rDNA sequencing from stool samples. Alpha-diversity was analyzed among the different subgroups of control mice. Richness (Observed feature) was significantly decreased in mice fed an ID or IE diet compared to mice fed an IA diet (means: 236.6 vs 281.3, *p-*value =2e-05 and 246.6 vs 281.3, *p* =4e-04 respectively, Figure 6A). However, diversity differences (Shannon index) between mice subgroups were not significant, except for a trend toward a decrease in mice fed an ID diet compared to mice fed an IE diet. Beta-diversity was evaluated by calculating distances between samples using Jaccard and Bray-Curtis methods. Principal coordinates analysis (PcoA) was performed to represent the distribution of biodiversity at the OTU level between groups (Figure 6B). Subgroups of mice fed an ID, IA or IE were clustered and significantly different (*p* <0.0001). We then examined the bacterial communities impacted by the dietary iron deprivation or supplementation. Relative abundance was analyzed at the phylum level (Figure 6C). A decreased proportion of *Bacillota* (previously known as *Firmicutes*) was observed in mice fed an ID (19%) or IE (10%) diet compared to mice fed an IA diet wherease an increase in the *Bacteroidota* (ID: 16%; IE: 4%) and *Pseudomonadota* (previously known as *Proteobacteria*) (ID: 4%; IE: 2%) was found. At the family level (Figure 6D), *Erysipelotrichaceae* (Bacillota phylum) was the most abundant family in mice fed an IA or IE diet, whereas *Tannerellaceae* (Bacteroidota phylum) was the predominant family in mice fed an ID diet. *Bacteroidaceae*, *Desulfovinrionaceae*, *Oscillospiraceae, Marinifilaceae* and *Tannerellaceae* families were more abundant in mice fed an ID or IE diet when compared to mice fed an IA diet. In contrast, *Akkermansiaceae*, *Erysipelotrichaceae*, and *Rikenellaceae* families were less represented in the microbiota of dietary challenged mice. However, an increase in *Muribaculaceae* family was observed only in mice with iron supplementation. Lastly, we assessed the significant changes in microbiota composition at the genus level depending on the dietary iron content using DESeq2. We observed a significant decrease or increase of 47 to 76 OTUs in mice fed an IE or ID diet compared to mice fed an IA diet (Figure 6E and supplementary material). In comparison to mice fed an IA diet, mice fed an ID or IE diet exhibited a significantly decreased abundance in *Akkermansia, Candidatus Saccharimonas, Faecalibaculum* and *Rikenellaceae RC9* and an increase in *Escherichia Shigella* with a more pronounced alteration in mice fed an ID diet (Figure 6F). *Parabacteroides* was the most abundant genus during iron deprivation along with a significant increase in *Bacteroides* and a reduction in *Alloprevotella* whereas mice fed an IE diet showed an increased abundance in *Alistipes* and *Blautia*.

**Figure 6:**
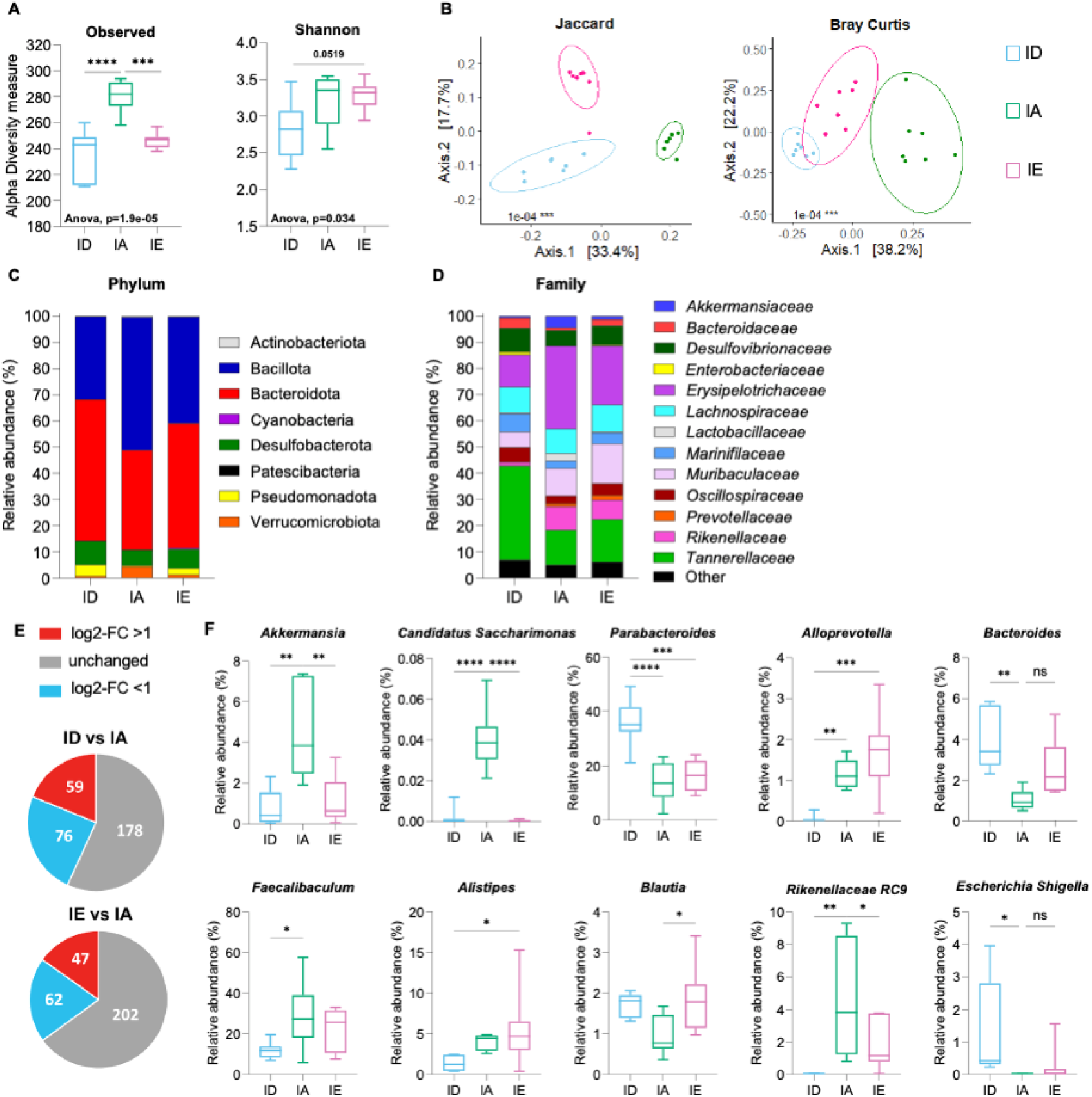
Dietary iron impact on fecal microbiota diversity. Assessment of the microbiome composition by 16S DNA sequencing from stool samples of control mice. (A) Amplicon sequence variants (ASV) distribution (alpha diversity): the observed number of species in each sample (richness) and Shannon (richness and evenness). (B) Principal component analysis of the similarities in microbial communities between groups (beta diversity) based on the Jaccard algorithm and Bray Curtis dissimilarity. (C) Phylum and (D) bacteria family. (E) Comparison of OTU levels depending on the dietary iron content. (G) Relative abundance of bacterial genus. Data shown are means ± s.e.m and were compared between DSS-treated mice and control mice for each diet (n = 8) by One-way ANOVA and corrected for multiple comparisons by Bonferroni method. **P < 0.01, *P < 0.05.

## DISCUSSION

Iron deficiency anemia is a common extra-intestinal complication affecting patients with inflammatory bowel diseases. The outcome of iron deficiency or iron supplementation in IBD has been extensively documented in mouse models and patients(19, 20). However, most animal studies have been performed using dietary iron restriction or supplementation that were not necessarily translatable to human studies. While common standard rodent chows contain 150-300 µg/g of iron, 20-25 µg/g of iron in the diet is sufficient to meet the daily iron needs in mice. In some studies, the iron deficient diet exceeded these daily requirements(9, 21–23) whereas a standard rodent chow was considered an iron enrichment. A 10-fold increase in enteral iron was reported to be protective whereas a 100-fold increase exacerbated the colitis in mouse(9, 11, 24, 25). In the present study, mice were fed an iron deficient diet (ID) containing 10 µg/g Fe, the minimum achievable considering the amount of iron found in cereals and other components of the diet. The impact of the dietary iron was compared to the same diet enriched with carbonyl-iron at a concentration of 50 µg/g (iron-adequate-IA) or 8000 µg/g (iron-enriched diet-IE) to match the oral iron supplementation administered to patients with iron-deficiency anemia. The dietary iron challenge allowed the accumulation of iron in the lumen of mice fed and IE diet but mice fed an ID or IE diet did not develop an iron deficiency or an iron overload in comparison to mice fed an IA diet (Figure 1).

To study the impact of dietary iron during colitis, mice were subjected to a murine model of chronic intestinal inflammation mimicking the flare-ups and recovery phases observed in patients which is based on two repeated cycles of 2% DSS or normal drinking water for 7 days each(26). However, after 9 days, mice fed an ID or IE diet had reached the maximal 20% body weight loss authorized by our ethical guidelines, which prompted us to compare the impact of dietary iron on the severity of colitis after 9 days (7 days 2% DSS, 2 days water). The impact of dietary iron deprivation or supplementation in a chronic DSS model could therefore not be tested. Of note, mice fed an ID diet exhibited earlier signs of gastrointestinal bleeding compared to mice fed an IE or IA diet suggesting an increased susceptibility to epithelial and vascular damage. Indeed, iron is necessary for the proliferation of stem cells during the regeneration that follows a tissue injury induced by DSS(27).

We did not detect major differences in mRNA expression of tight junction proteins, inflammatory mediators and markers of ferroptosis and oxidative stress between mice fed an ID and IE diet except for a higher expression of *Tnfa* and *Il17a* in the colon of DSS-treated mice fed an IE diet. These results suggest that the presence of iron in the lumen does not directly affect the expression of tight junction proteins. A potential explanation for the increase in *Tnfa* expression is that iron induces the polarization of proinflammatory macrophage (M1) through the mitogen-activated protein kinase (MAPK) pathway(28). Our results also suggest a similar degree of oxidative stress and ferroptosis in mice fed an ID or IE diet (Figure 5). Both groups of mice showed early signs of bleeding and, while heme iron is not detectable by Perls’ staining, heme iron may accumulate in the colonic lumen leading to oxidative stress, ferroptosis and epithelial damage(29), independently of the dietary iron content.

The amount of iron in the diet is known to alter the microbial composition in the colon by exerting a selective pressure or by promoting the proliferation of specific bacteria(30). While a different dysbiosis has been reported depending on the iron formulation used to supplement the diet(11), our results show that alterations in dietary content led to a profound remodeling of bacterial composition and an increased susceptibility to colitis. Indeed, the alpha- and beta-diversity analyses revealed a dysbiotic state of the fecal microbiota of mice fed an IE or ID diet compared to mice fed an IA diet. In addition to its essential roles in the host and microorganisms, iron disturbs the equilibrium of the microbial composition by promoting the growth and pathogenesis of siderophilic bacteria(30). Interestingly, dietary iron deprivation and supplementation resulted in similar changes in the most abundant gut microbial phyla represented by a decreased proportion of Bacillota and an increase in Bacteriodetes and Pseudomonadota. The phylum Verrucomicrobiota was less abundant in mice fed an ID diet in comparison to mice fed an IA diet. Studies in IBD patients also reported a reduced level of short-chain fatty acid producers Bacillota and a pronounced abundance of Pseudomonadota(31, 32). Among the Bacillota, the abundance of the Erysipelotrichaceae family was reduced in mice fed an IE or ID diet similar to CD patients(33). This family includes *Faecalibaculum,* a bacterium known to influence regulatory T cells and reduce colonic inflammation(34). Similarly, manipulating dietary iron resulted in a decreased abundance of *Akkermancia*, a promising probiotic against colitis that is also reduced in IBD patients(35). The most notable change in the microbial family observed only in mice fed an ID diet was the increased abundance of *Tannerellaceae* and specifically *Parabacteroides*, which have been shown to reduce intestinal and systemic inflammation and obesity(36).

Pseudomonadota phylum includes pathogenic microorganisms such as the *Enterobacteriaceae* family and the well-known *Escherichia*. In the present study, we found a greater genus variation in *Escherichia Shigella* between mice fed an ID or IE diet compared to mice fed an adequate chow. These siderophilic gram-negative species are responsible for enteric infections worldwide, as a result of the invasion and destruction of the colonic mucosa that leads to acute diarrhea, hemorrhage and ulceration(37). However, the physiological relevance of *E. Shigella* in murine models is still debated(38).

The rapid change (9 days) in microbiota composition in mice fed an ID diet may not be found in humans eating an omnivore diet with sufficient iron intake. Indeed, only 10% of the iron present in the food is absorbed in the duodenum while 90% transits through the intestinal tract. A systemic iron deficiency caused by hepcidin-mediated iron restriction would not lower the amount of iron found in the lumen unless the patients consume only iron-poor products. However, as suggested in previous studies; this work confirms that iron supplementation can fuel the inflammation during IBD, thus questioning the benefit/risk balance of oral iron supplementation for IBD patients. As iron absorption is impaired in patients with Crohn’s disease, iron deficiency can be corrected with intravenous iron formulations to bypass the gastrointestinal tract absorption but the inconvenience of IV administration is often accompanied by a broad range of side effects(7).

## ACKNOWLEDGMENT

The authors thank Jorge Ayala and Manon Sotin for technical assistance, members of the INSERM US006 animal and histopathology facilities (Toulouse) and the platform Aninfimip, an EquipEx (‘Equipement d’Excellence’) supported by the French government through the Investments for the Future program. We are grateful to the INRAE MIGALE bioinformatics facility (MIGALE, INRAE, 2020. Migale bioinformatics Facility, doi: 10.15454/1.5572390655343293E12) for providing computing and storage resources and to the Metagenomic16s service for microbiota analyses (CRI, Paris, France). Support for this work was provided by the French National Research Agency (ANR-16-ACHN-0002-01 and ANR-22-CE14-0076-01), the “Fondation Maladie Rare”, the region Occitanie and by the European Research Council (ERC) under the European Union’s Horizon 2020 research and innovation program (grant agreement no. 715491) to LK, The French Society for Hematology (SFH) to US.

## Authorship Contributions

TM and AD performed experiments, analyzed data and wrote the paper. US, PP, KC, MR performed experiments, LK designed and supervised the study, analyzed data and wrote the manuscript. All authors edited the manuscript.

